# Attentional modulation of neural phase is enhanced by short-term training and linked to musical experience

**DOI:** 10.1101/519181

**Authors:** Aeron Laffere, Fred Dick, Adam Tierney

## Abstract

How does the brain follow a sound that is mixed with others in a noisy environment? A possible strategy is to allocate attention to task-relevant time intervals while suppressing irrelevant intervals - a strategy that could be implemented by aligning neural modulations with critical moments in time. Here we tested whether selective attention to non-verbal sound streams is linked to shifts in the timing of attentional modulations of EEG activity, and investigated whether this neural mechanism can be enhanced by short-term training and musical experience. Participants performed a memory task on a target auditory stream presented at 4 Hz while ignoring a distractor auditory stream also presented at 4 Hz, but with a 180-degree shift in phase. The two attention conditions were linked to a roughly 180-degree shift in phase in the EEG signal at 4 Hz. Moreover, there was a strong relationship between performance on the 1-back task and the timing of the EEG modulation with respect to the attended band. EEG modulation timing was also enhanced after several days of training on the selective attention task and enhanced in experienced musicians. These results support the hypothesis that modulation of neural timing facilitates attention to particular moments in time and indicate that phase timing is a robust and reliable marker of individual differences in auditory attention. Moreover, these results suggest that nonverbal selective attention can be enhanced in the short term by only a few hours of practice and in the long term by years of musical training.

## Introduction

Humans are constantly bombarded by multiple streams of information, some vitally important, but most not. One of the central computational problems facing the brain is how to extract the most relevant signals while filtering out irrelevant information and noise. This problem is particularly difficult for the auditory system which - unlike the visual system - cannot rely on mechanical means (saccades, focusing) to screen out unwanted information. Whether listening to a conversational partner in a crowded pub, or focusing on the bassist in a jazz ensemble, listeners must somehow separate out the different auditory streams (auditory object formation) and devote in-depth processing to one of those streams (auditory object selection (1)).

When task-relevant auditory objects have characteristic temporal regularities, attention to time can be used to select target objects and suppress distractor objects (2). That attention can be directed towards moments in time is supported by research showing that participants produce faster responses when stimuli appear at expected times. This has been demonstrated for a variety of perceptual tasks in both the auditory (3–4) and visual (5–6) systems. Attention to time can not only be used to enhance performance in single-stream perception, but can also be used to facilitate object selection. Prior knowledge about auditory stream timing facilitates listening in cocktail party paradigms (7–9), while uncertainty about stimulus timing makes tone detection more difficult when distractor tones are also present (10). Moreover, differences in speech rate facilitate perception of speech presented alongside distracting speech (11), and talkers producing speech alongside distractor talkers will adjust their speech rate to reduce temporal overlap with the competing speech (12). One strategy, therefore, which listeners could use to facilitate auditory object selection is that attention could be directed towards points in time when target objects are likely to appear, and away from points in time when distractor objects are likely to appear (13). For example, if a listener knows her conversational partner well, she could use her knowledge of the talker’s speaking rate to predict when the next word will begin, and focus attention on that time point.

Over the last decade cognitive neuroscience research on speech perception has uncovered a possible neural mechanism underlying auditory object selection based on temporal information. Neural activity synchronizes to low-frequency fluctuations in the amplitude of speech, as measured using electroencephalography (EEG) and magnetoencephalography (MEG) (14–24). Selecting one auditory stream and ignoring another could therefore involve enhancing neural entrainment to the rhythmic patterns in the target stream, and diminishing entrainment to the patterns in the distractor stream (25). Indeed, when multiple speech streams are presented, neural entrainment to the temporal envelope of both the target and the background is present, but that of the target speaker is enhanced relative to the distractor speakers (15, 26–29), and the extent of this modulation correlates with speech comprehension scores (30). Moreover, transcranial stimulation with currents shaped similarly to the amplitude envelope of target speech can facilitate speech recognition performance in cocktail party paradigms and fMRI responses to speech (24, 31), demonstrating that neural entrainment plays a causal role in auditory object selection.

Research on neural entrainment and selective auditory attention in humans has largely focused on speech perception. Comparatively little, therefore, is known about the neural mechanisms underlying more general (non-verbal) selective auditory attention. Neural entrainment is not limited to speech stimuli, and has been demonstrated to simple abstract sounds (32–33), non-verbal rhythms (34), and ecologically valid music (35–38) as well. Research on non-human primates in which non-verbal stimulus streams are presented at different time points has shown that switching attention between streams modulates neural entrainment (39–42). However, there is almost no prior research on neural entrainment and non-verbal selective auditory attention in humans. The only exceptions, to our knowledge, are Besle et al. (43), who demonstrated attentional modulation of neural entrainment when human epilepsy patients were asked to attend to either an auditory or a visual stimulus stream, and Morillon and Baillet (44), who demonstrated modulation of stimulus-response phase-locking when participants are asked to attend to and judge the pitch of tones presented in alignment with a beat, and ignore tones out of alignment with the beat. Other EEG research on selective attention to a target melody has analyzed attentional modulation of onset responses rather than neural entrainment (45–46).

Despite the importance of robust auditory selective attention for everyday listening, there are large individual differences in the ability to listen and respond to single auditory streams in the presence of distractors (47). The neural foundations of individual differences in selective attention are however poorly understood, as previous studies of the neural correlates of selective attention have used small numbers of participants and focused their analyses on main effects of attention. Here we hypothesized that individuals who are skilled at directing attention to an auditory object show stronger top-down modulation of neural entrainment to sound. We tested this hypothesis by asking participants to respond to targets in an attended sequence of tones while suppressing attention to a competing sequence in a different frequency band. The sequences were presented at the same rate but interleaved so that distractor tones were in counter-phase with targets. We predicted that active attention would be linked to neural entrainment to the attended sequence, leading to an increase in inter-trial phase locking at the frequency of within-band stimulus presentation (4 Hz). Moreover, we predicted that the direction of auditory selective attention would be linked to an approximately 180-degree shift in the phase of the EEG signal at 4 Hz, and that the extent of this shift would correlate with task performance.

Prior studies comparing cognitive abilities in musicians and non-musicians have found that musicians display an advantage for selective attention in both visual (48) and auditory (49–56; but see 57–58) stimuli. This suggests that the neural mechanisms underlying selective attention may be responsive to training. However, the musicians in these studies experienced thousands of hours of training, beginning in childhood, and so it remains unclear whether enhancement in selective attention is possible after shorter-term training in adulthood. Moreover, the neural foundations of training-based selective auditory attention enhancements have not yet been investigated. Given that musicians show increased cortical entrainment to music (36,38), one possibility is that musicians show stronger top-down modulation of neural entrainment to sound due to their experience directing attention to a single sound source within a highly complex acoustic environment. Here we tested this hypothesis by examining inter-trial phase locking and average neural phase during auditory selective attention in participants with and without musical training. We predicted that musicians would show greater neural phase differentiation between attention conditions, as well as increased inter-trial phase locking at the within-band stimulus presentation frequency. Finally, to examine the effects of short-term training on attentional modulation of neural entrainment and auditory selective attention ability, we tested participants before and after a two hour online attention training session. We predicted that online training would lead to enhanced performance, as well as enhanced inter-trial phase locking and neural differentiation of attention conditions.

## Results

### Task overview

Participants (*n* = 43) were asked to attend to tones in a low (185 - 233 Hz) or high (370 - 466 Hz) frequency band while ignoring tones presented concurrently in the competing frequency band (see Fig. 1 for a task schematic). In a third “Passive” condition participants sat quietly and listened to the tones but were not asked to actively attend to either band. Tones in contrasting frequency bands were asynchronous and formed a repeating ABABAB pattern of low (A) and high (B) frequencies followed by a silent pause at the end of each mini-sequence. For each band, a set of three possible fundamental frequencies was randomly sampled to create mini-sequences with a length of three tones. If a mini-sequence was repeated consecutively in the attended frequency band, the participant was asked to respond as quickly as possible by clicking the mouse. Tones were presented 180 degrees out of phase at regular intervals with a within-sequence rate of presentation of 4 Hz (i.e. 250 ms between tone onsets) and a cross-sequence rate of presentation of 8 Hz (125 ms between tone onsets).

**Figure 1.**
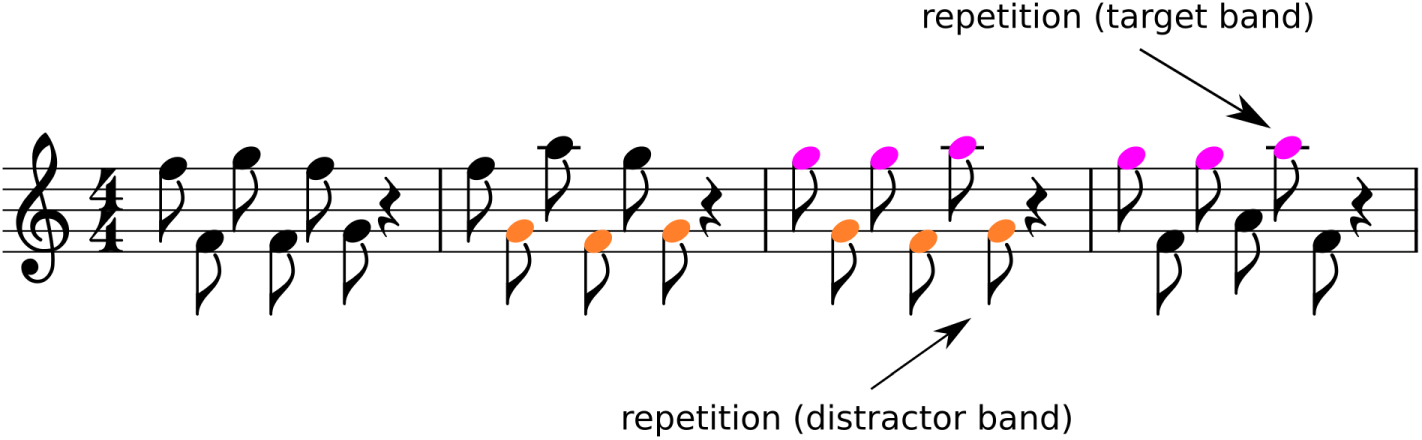
Schematic of non-verbal selective attention task. Participants were asked to listen to short melodies made up of three tones in a target frequency band and press a button whenever they heard a repeated sequence, while ignoring repetitions in a distractor frequency band. The target and distractor sequences were interleaved, such that when a tone from one sequence was present the other sequence was silent.

### Channel selection

To limit the analysis to channels where entrainment was most strongly present, any EEG channel in which intertrial phase coherence was less than 0.1 at 4 Hz averaged across all conditions was excluded from further analyses (of 32 original channels, 14 channels remained: Fp1, AF3, F3, FC1, FC5, C3, CP1, C4, FC6, FC2, F4, AF4, Fz, and Cz).

### Effects of selective attention on neural phase and inter-trial phase coherence

To examine whether actively directing attention to one of the two frequency bands affected the strength of neural entrainment, we calculated inter-trial phase coherence (ITPC) at 4 Hz (the within-band presentation rate). ITPC was higher for the active conditions (i.e. attend high and attend low, M = 0.115, SD = 0.053) than for the passive condition (M = 0.092, SD = 0.052; t (42) = 2.15, p < 0.05). We also calculated ITPC at the cross-frequency-sequence rate of stimulus presentation (8 Hz) and found no significant difference between active and passive conditions (t(42) = 0.14, p > 0.1), suggesting that increased neural entrainment was specific to the within-band repetition rate.

Next, to test the hypothesis that the direction of auditory selective attention would be linked to a shift in the phase of the EEG signal at 4 Hz, we investigated whether the direction of attention (i.e. attend high or attend low) led to a change in the average neural phase across conditions. We used a Watson-Williams test for equal means in circular data, as implemented in the circular statistics toolbox for Matlab (59). This test could not be conducted on the entire dataset because not all assumptions were met, due to the average resultant vector being < 0.45. Given the high degree of variability in performance across participants - hit rates ranged from 11.1% to 82.5% of repetitions during the first testing session (M = 48.7%, SD = 14.2%) - we repeated this analysis on a dataset limited to the half of the participants who performed best on the task (n = 22). The test could be run on this smaller dataset, and as predicted above there was a significant difference between the average phase angles at 4 Hz for the attend high (M = 2.60 radians) and attend low (M = −1.07 radians) conditions (F(1,44) = 19.01, p < 0.0001), indicating a modulation of neural phase by the direction of selective attention in participants who more successfully performed the behavioral task.

### Correlations between behavioral performance and neural metrics

Given the high degree of performance variability, we investigated correlations between behavioral performance and several neural metrics. First, we investigated the relationship between performance and strength of neural entrainment by calculating ITPC. Task performance was not associated with ITPC at the within-band presentation rate (4 Hz) during the active conditions (R^2^ = 0.08, p > 0.05). Next, we investigated whether particular average neural phases at 4Hz were linked to better performance. Each participant’s average neural phase was converted to Cartesian coordinates, and these two coordinates were included as predictors in a linear regression, with hit rate across all conditions as the dependent variable. There was a significant relationship between performance and average neural phase in attend high (R^2^ = 0.16, p < 0.05) and attend low (R^2^ = 0.22, p < 0.01) conditions. The regression equation was then used to extract behaviorally optimal neural phases, i.e. phases which were linked to the highest hit rates. This was done by using the regression equations to calculate predicted performance for each possible neural phase, then selecting the neural phase that maximized predicted performance. Behaviorally optimal phases were roughly π radians apart across the two attention conditions: 2.2531 radians for attend high and −0.9837 radians for attend low, for a difference between behaviorally optimal phases of 3.0464 radians. For the subsequent analyses, distance from behaviorally optimal phase is calculated as the distance between the individual's average neural phase in each attention condition and the behaviorally optimal phase for that condition. Collapsing across both attention conditions, there was a significant negative correlation between performance and distance from behaviorally optimal neural phase at 4 Hz (Spearman’s correlation, R^2^ = −0.41, p < 0.0001; Fig. 2B). In Fig. 2A participants are separated by using the same median split based on performance as in the phase analysis above to illustrate the relationship between neural phase and performance.

**Figure 2.**
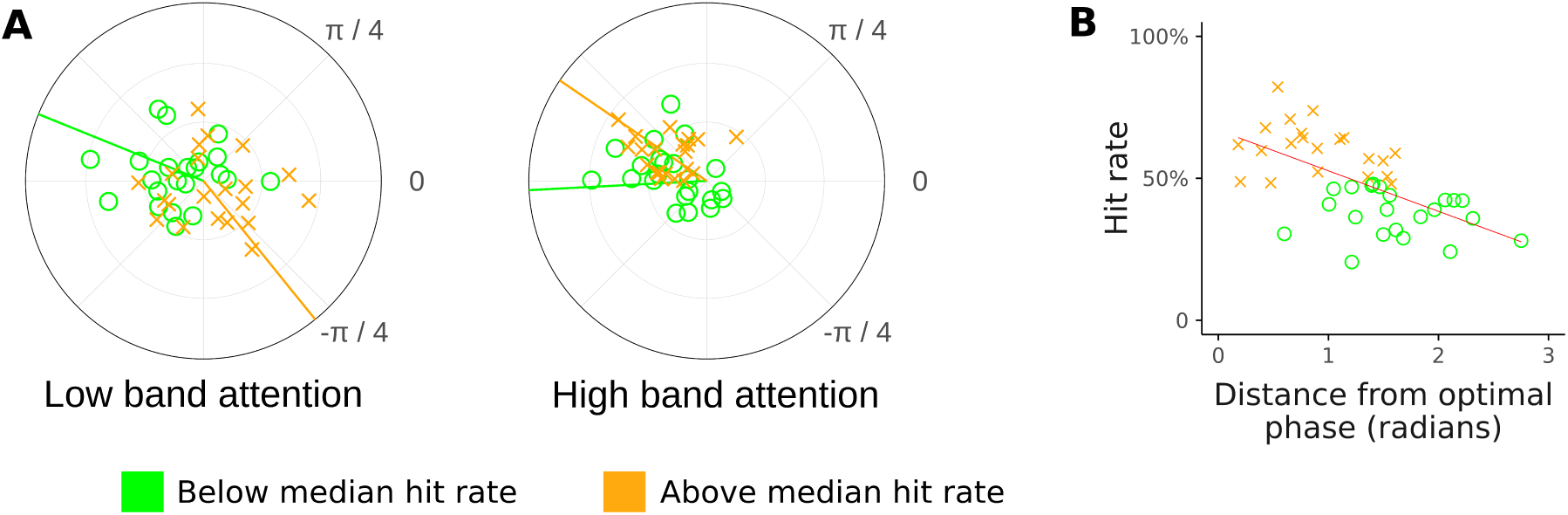
(A) Average neural phase at the within-band presentation rate in attend high and attend low conditions in participants with low (green circles) versus high (orange crosses) hit rates (median split). Data points represent both inter-trial phase consistency (distance from origin) and average phase (angle). Lines represent the average neural phase across participants for each group. (B) Scatterplot displaying the relationship between hit rate (detection of repeated sequences) and behaviorally optimal neural phase. Optimal phase was extracted by regressing neural phase versus performance.

### Group differences between musicians and non-musicians

We defined musicians (*n* = 18) as participants with three or more years of training in a musical instrument and non-musicians (*n* = 18) as participants with no musical training. (7 participants who did not fit into either category were excluded from the analysis.) Average neural phase and ITPC for musicians and non-musicians is illustrated in Fig. 3A and B. More targets were detected by musicians than by non-musicians (t(34) = 3.74, p < 0.001; Fig. 3C). There was no significant difference in ITPC between musicians and nonmusicians at 4 Hz in either active (t(34) = 0.01, p > 0.1) or passive (t(34) = −1.35, p > 0.1) conditions, nor at 8 Hz in either active (t(34) = 1.56, p > 0.1) or passive (t(34) = −0.07, p > 0.1) conditions. However, musicians' behavioral advantage was accompanied by more precise neural modulation by attention, as the distance from behaviorally optimal neural phase at 4 Hz was smaller for musicians than for non-musicians (t(34) = 2.91, p < 0.01).

**Figure 3.**
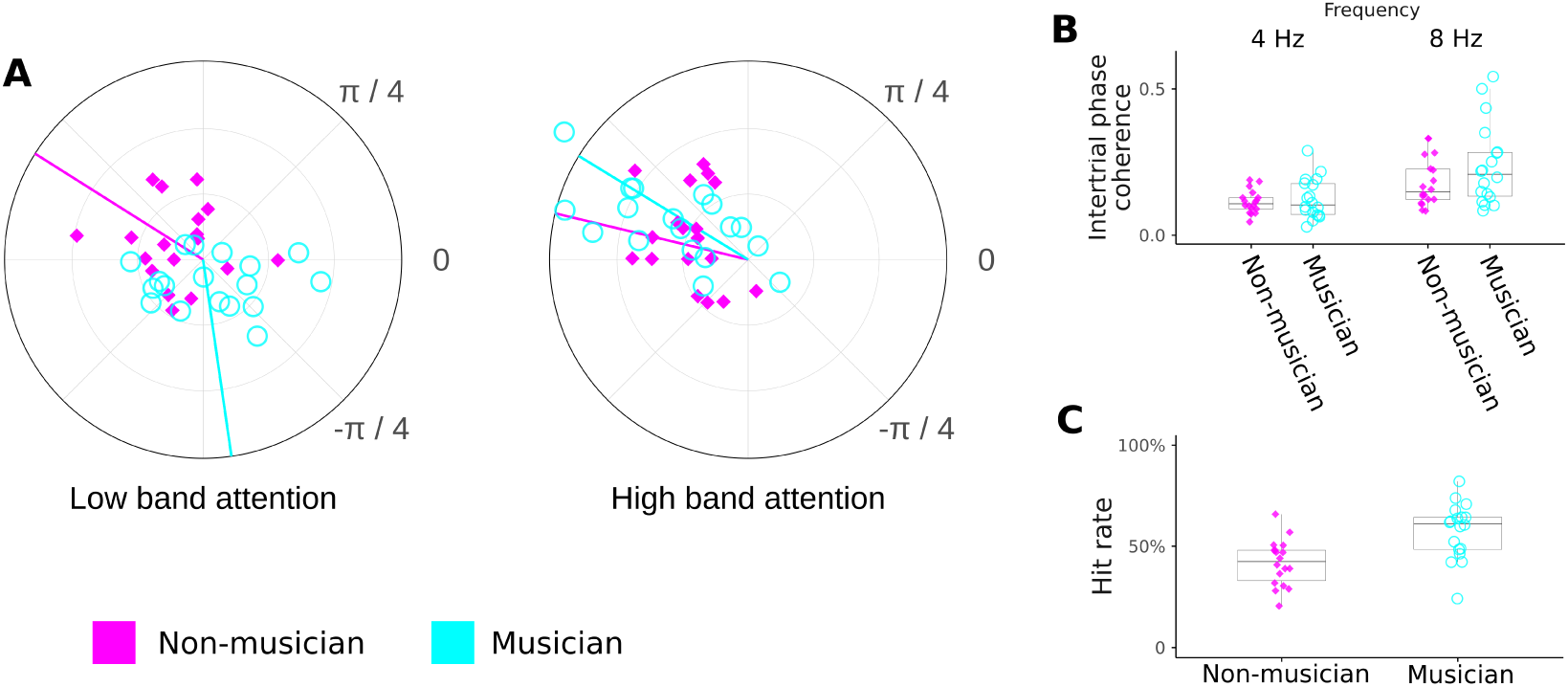
(A) Average neural phase at the within-band presentation rate in attend high and attend low conditions in musicians (blue circles) and non-musicians (pink diamonds). (B) Inter-trial phase coherence at within-band and between-band presentation rates in musicians and non-musicians. (C) Rate of target detection for musicians and non-musicians.

### Within-subjects analysis of effects of online training

Participants (*n* = 24) who completed a two-hour online training task and returned for a second EEG recording session performed significantly better on the task after training (t = 10.92, p < 0.0001; Fig. 4A and B). There was no significant effect of training on intertrial phase coherence at 4 Hz in active conditions (t = 1.27, p > 0.1; Fig. 4C and D). By contrast, distance from behaviorally optimal neural phase at 4 Hz was smaller after training than before training (paired t-test, t(23) = −3.55, p = < 0.01, Fig. 4E and F). Fig. 5A and B illustrate the hit rate and distribution of phase angles across participants before and after the training session.

**Figure 4.**
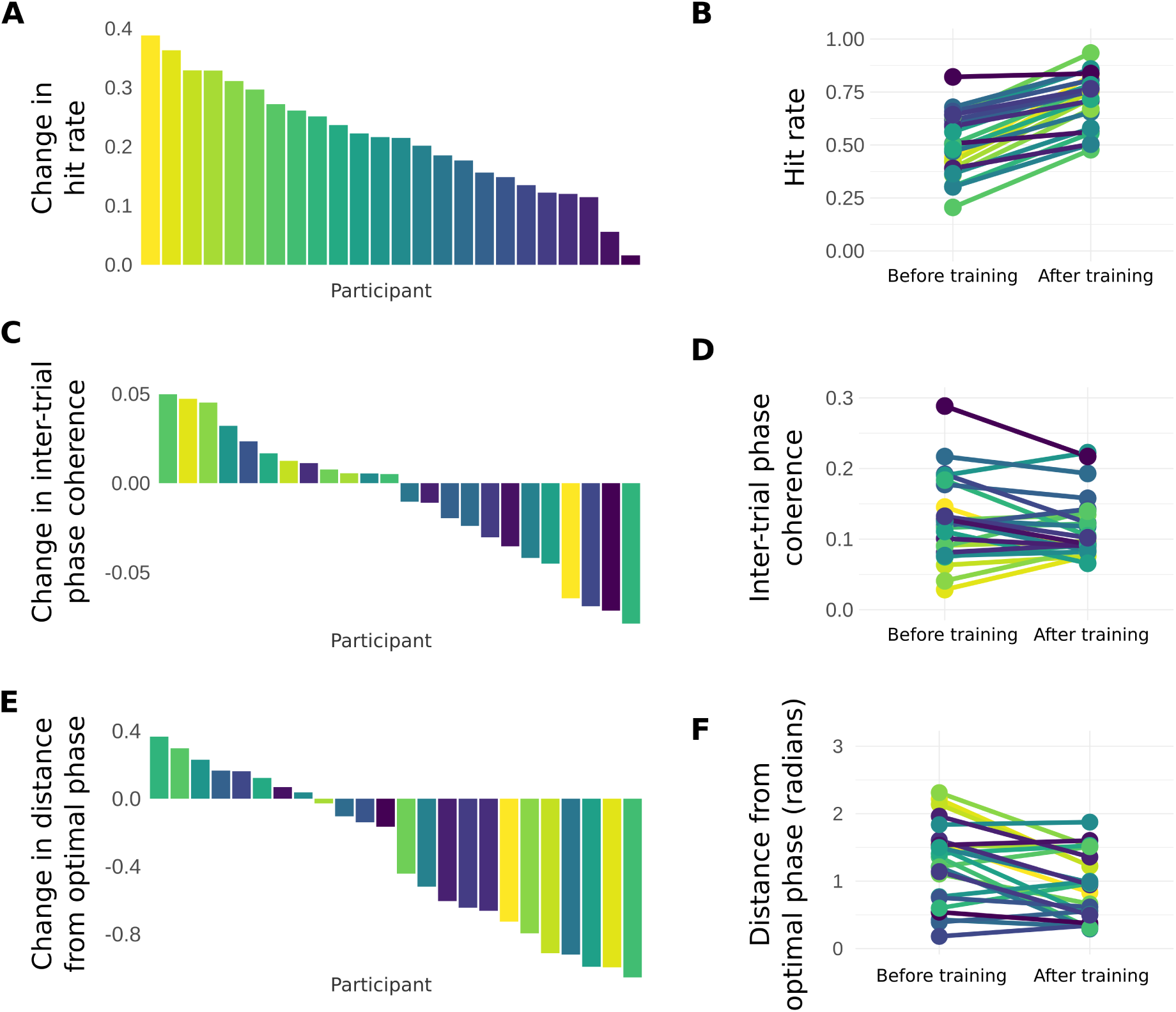
(A) Change in hit rate between pre-training and post-training testing sessions. (B) Hit rates in pre-training and post-training sessions. (C) Change in inter-trial phase coherence between pre-training and post-training sessions. (D) Inter-trial phase coherence in pre-training and post-training sessions. (E) Change in distance from optimal neural phase in pre-training and post-training sessions. (F) Distance from optimal neural phase in pre-training and post-training sessions.

**Figure 5.**
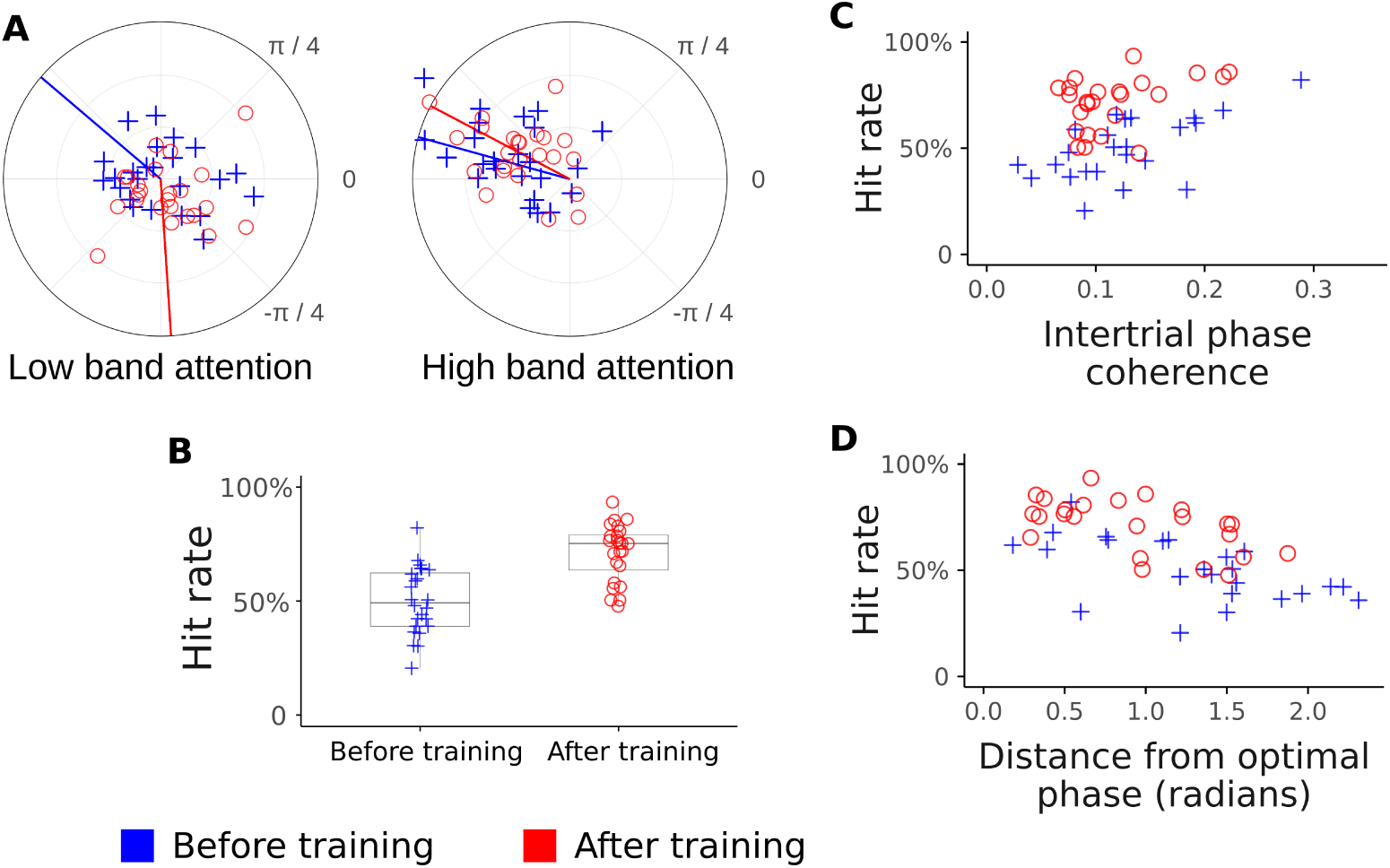
(A) Average neural phase at the within-band presentation rate in attend high and attend low conditions in pre-training (blue crosses) and post-training (red circles) testing sessions. (B) Hit rate in pre-training and post-training sessions. (C) Relationship between inter-trial phase coherence and hit rate in pre-training and post-training sessions. (D) Relationship between distance from optimal neural phase and hit rate in pre-training and post-training sessions.

## Discussion

Here we report a new phase-based method for tracking selective attention to non-verbal sounds using EEG, and show that this neural metric is closely tied to performance, enhanced in musicians, and boosted by short-term training. We find that selective attention, compared to passive listening, is linked to an increase in inter-trial phase coherence at the within-band presentation rate. Moreover, we find that attention to one of two alternating stimulus streams is linked to a 180-degree offset in neural phase. This suggests that one of the ways listeners can select a target stream for in-depth processing is by directing attention to moments of time at which a target is likely to appear (and away from moments in time at which a distractor is likely to appear), and that this temporally-directed attention is linked to phase alignment of neural rhythms with the attended stimulus stream. This finding is consistent with previous reports of electrophysiological studies in non-human animals and electrocorticographical studies of human epilepsy patients indicating that switching attention from one non-verbal stimulus stream to another is linked to a shift in neural phase (39–43).

We show that an individual's average neural phase can be used as a metric of inter-subject differences in selective attention. Moreover, participants’ neural responses were more tightly clustered around this behaviorally optimal phase after training, and musical participants who had undergone musical training had neural responses that were closer to this behaviorally optimal phase. Thus, the optimal phase relationship between neural activity and stimulus timing is surprisingly consistent across participants, and can be used to assess individual differences in selective attention. By comparison, inter-trial phase coherence was less consistently linked to performance, was not modulated by training, and was not related to degree of musical training. These results suggest that the extent to which participants are able to align neural activity with the target stimulus stream is one factor underlying individual differences in selective attention ability, while the consistency of neural entrainment may be less important.

Compared to participants with no musical training, those with at least three years of musical training showed more precise alignment between their neural response and the attended stimulus, as well as a sizeable advantage in performance, suggesting that musical experience can enhance selective attention to sound and top-down modulation of neural entrainment. However, we found that inter-trial phase coherence did not differ between musicians and non-musicians in any condition, whether at the within-band or the across-band presentation rate. This suggests that the musician advantage was specifically for attentional modulation of neural phase, rather than a more general enhancement of neural entrainment regardless of task. These results are somewhat inconsistent with previous reports that participants with musical training show enhanced cortical responses even in passive listening paradigms (60–65) and increased cortical entrainment to music (36,38). Our lack of a musician advantage for inter-trial phase locking may reflect the simplistic nature of the stimuli: it is possible that musicians may demonstrate enhanced entrainment only for stimuli which are timbrally, melodically, or rhythmically complex. However, these results are consistent with prior reports that musicians display an advantage for selective attention in visual (48), verbal auditory (50-56; but see 57-58), and non-verbal auditory (49) stimuli.

We found that only a few hours of training led to a considerable increase in selective attention performance, with an average performance gain of 21% between the pre-training and post-training testing session. Moreover, we found that training also led to an increase in the alignment of neural phase with the attended sound stream, suggesting that individuals can rapidly improve their top-down control over neural entrainment. This concomitant neural enhancement demonstrates that the performance improvement was at least partially due to improved attentional control, rather than simply to an increased ability to discriminate melodic patterns. These behavioral and neural enhancements demonstrate the possibility of rapid short-term plasticity in the mechanisms of selective auditory attention, suggesting that short-term training programs may be a successful remediation strategy in populations who struggle to control attention.

Our result suggest several potentially fruitful lines of fruitful work. For example, our phase-based analysis cannot distinguish between contributions from neural enhancement of the attended stimulus stream and neural suppression of the ignored stimulus stream (13). Future work could distinguish between these two possibilities by presenting the two streams at different rates—for example, 4 Hz and 3 Hz—to determine whether neural entrainment to a given stream is enhanced when it is attended and suppressed when it is ignored relative to a passive condition. It also remains unclear whether musicians’ enhanced top-down modification of neural entrainment is limited to musical stimuli or whether it extends to other domains as well. This could be tested by examining attentional modulation of neural entrainment to the speech envelope in musicians and nonmusicians. Similarly, future studies could investigate whether the effects of the non-verbal attentional training paradigm introduced in the current study generalize to behavioral and neural measures of attention to speech. Finally, because we did not include control training in our design, we cannot draw any conclusions about the efficacy of this particular selective attention training program relative to any alternate form of training, and future work could use this paradigm to examine the factors which modulate the efficacy of attention training programs. Moreover, future work should examine the extent to which selective attention enhancements are maintained long-term if training is discontinued.

In conclusion, we demonstrate that selective auditory attention is linked to a modulation of neural phase and that this metric is tightly linked to individual differences in selective attention performance. Our results further suggest that attentional control, as measured both behaviorally and neurally, can be enhanced by both short-term training and long-term experience.

## Materials and Methods

### Participants

The subjects in this experiment were recruited over the course of several months from a population of adults with no history of neurological disorder or hearing impairment. All study procedures including informed consent and incentives were approved by the Birkbeck Department of Psychological Sciences ethics committee. 43 subjects (ages 18 - 43 years, M = 28.9, SD = 7.5, 23 female) participated in the first testing session. After any subject who failed to complete training or missed their second appointment was excluded from the study, twenty-four subjects (ages 18 - 43 years, M = 28.6, SD = 7.5, 11 female) remained for the second session. Hearing thresholds were assessed prior to study procedures by presenting tones at audiometric frequencies from 500 to 4000 Hz. Any subject who failed to hear tones presented at 20 dB SPL in a pretest assessment was excluded from analyses.

### Procedure

Participants were first tested in a two-hour pre-training session in which they were asked to selectively attend to tone sequences in a target frequency band, detecting occasional repeats, while ignoring tone sequences in a distractor frequency band. Feedback for correct responses, missed targets, and incorrect responses was presented, and EEG data was concurrently collected. On the next day participants completed a two-hour online training session in which they completed the same task with similar feedback. On the third day participants returned to the lab to complete a two-hour post-training EEG session during which they completed the same task.

### Stimulus design and presentation

The fundamental unit of stimulus presentation was a 125 ms cosine-ramped pure tone generated at a sampling rate of 48 kHz using Matlab. A trial consisted of six tones presented without pause (750 ms) followed by a silence lasting the duration of two tones (250 ms) for a total trial length of 1 s. Tones were presented within two frequency bands. Within each frequency band tones were drawn from a set of three tones, each two semitones apart. For the low band, tones could be set to frequencies of 185, 207.7, and 233.1 Hz; for the high band, tones could be set to frequencies of 370, 415.3, and 466.2 Hz. Tones in the high and low frequency bands alternated, such that a high frequency tone was always followed by a low frequency tone, and vice versa. Tones in the high frequency band were presented at 40% of the amplitude of tones in the low frequency band to ensure that the loudness of the two bands was relatively balanced. 3-6 within-band repetitions of the carrier frequencies drawn for two consecutive sequences were randomly distributed across each trial. Subjects were asked to detect and respond (with a computer mouse click) to within-band repetitions in the target frequency band defined for them by the experimenter while ignoring tones in the alternative frequency band. The latency window for a response to be recorded was from 250 ms before to 1250 ms after the end of the last tone in a sequence. Textual feedback was presented on a plain background computer display to notify subjects of correct responses, missed targets, and incorrect responses.

EEG recording sessions differed slightly in procedure from online training. EEG recording sessions consisted of thirty-five sets of thirty trials for each target frequency band and a third set for which subjects were asked not to respond to any repetitions. ER-3A insert earphones (Etymotic Research, Elk Grove Village, IL) were used for sound presentation at 80 dB SPL during these sessions. Online training began with 5 sets of thirty trials only containing tones from a target frequency band. Tones from the alternative frequency band were introduced in the 6th set at -30 dB and then increased in amplitude by 5 dB every subsequent 3 sets until reaching their original amplitude presentation. Subjects then completed seventy sets of thirty-five trials at the typical amplitude presentation before repeating the above procedure while targeting the other frequency band. As this part of the study was completed online, all sounds were presented through earphones available to the subject at a comfortable amplitude of their choosing. Aside from these differences, EEG recording sessions and training shared the feature that within-sequence repetitions appeared between three and six times in each set of thirty-five trials. After a repetition occurred, there was always at least one intervening non-repeating sequence before another repetition could begin. Trials in each set were presented without pause to maintain a steady presentation rate. Opportunities for rest were made available to subjects at the end of each set of trials. The full study procedure lasted around two hours on each day.

### EEG data acquisition and processing

All data were recorded from a BioSemi™ ActiveTwo 32-channel EEG system and digitized with 24-bit resolution. All channels were linked to Ag-AgCl active electrodes recording at a sampling rate of 16,384 Hz and positioned in a fitted headcap according to the standard 10/20 montage. External reference electrodes were placed at the ear lobes to record unlinked data for offline re-referencing. Contact impedance was maintained beneath 20 kΩ. Markers for the beginning of each set of 30 trials were recorded from trigger pulses sent to the neural data collection computer. The resultant data were downsampled to 500 Hz and then segmented into 1 s epochs aligned with trial onsets. Sources of artefact such as muscle noise and eye blinks were identified by independent component analysis of the epoched recording and removed after visual inspection of component topographies and time courses. Any epoch containing signal intensity in any individual channel exceeding +/- 100 µV was then rejected. Time-frequency analyses were carried out via Hann-windowed fast Fourier transforms. All pre-processing steps were carried out using Matlab and a blend of custom and premade scripts from the FieldTrip M/EEG analysis toolbox (66).

### Analyses

#### Behavioral data

The proportion of presented targets which were correctly identified was averaged across trials for each participant to generate performance measures for the high frequency and low frequency attention conditions. When determining median performance or making comparisons between groups or different time points, performance was averaged across attend high and attend low conditions.

#### Intertrial phase coherence

Intertrial phase coherence is a measure of the consistency or concentration of phase values across trials at a particular frequency or in a frequency band. Unlike measures of phase between channels which relate to connectivity, intertrial phase coherence measures the phase consistency across all trials separately in each channel. Phase coherence ranges from 0 (no consistency in phases whatsoever) to 1 (perfect phase alignment).

#### Statistical analyses

Statistical analysis of circular data was conducted using the Circular Statistics Toolbox (59). All analyses were conducted in Matlab using two-tailed p-values. Assumptions of normality for pairwise comparisons were checked and non-parametric test alternatives were used where appropriate. All data are available on request to the corresponding author.

## Acknowledgements

The authors are grateful to Maria Chait, Alex Billig, and Sijia Zhao for comments on an earlier version of the manuscript. None of the authors have potential conflicts of interest to disclose. The authors would also like to thank the participants for generously donating their time.

## References

1. Shinn-Cunningham B (2008) Object-based auditory and visual attention. Trends in Cognitive Sciences 12: 182–186.

2. Nobre A, van Ede F (2018) Anticipated moments: temporal structure in attention. Nature Reviews Neuroscience 19: 34–48.

3. Riess Jones M, Moynihan H, MacKenzie N, Puente J (2002) Temporal aspects of stimulus-driven attending in dynamic arrays. Psychological Science 13: 313–319.

4. McAuley J, Riess Jones M (2003) Modeling effects of rhythmic context on perceived duration: a comparison of interval and entrainment approaches to short-interval timing. Journal of Experimental Psychology: Human Perception and Performance 29: 1102–1125.

5. Lasley D, Cohn T (1981) Detection of a luminance increment: effect of temporal uncertainty. J. Opt. Soc. Am. 71: 845–850.

6. Rohenkohl G, Coull J, Nobre A (2011) Behavioral dissociation between exogenous and endogenous temporal orienting of attention. PLoS ONE 6: e14620.

7. Best V, Ozmeral E, Shinn-Cunningham S (2007) Visually-guided attention enhances target identification in a complex auditory scene. JARO 8: 294–304.

8. Gatehouse S, Akeroyd M (2008) The effects of cueing temporal and spatial attention on word recognition in a complex listening task in hearing-impaired listeners. Trends in Amplification 12: 145–161.

9. Kitterick P, Bailey P, Summerfield A (2010) Benefits of knowing who, where, and when in multi-talker listening. JASA 127: 2498–2508.

10. Bonino A, Leibold L (2008) The effect of signal-temporal uncertainty on detection in bursts of noise or a random-frequency complex. JASA 124: EL321–EL327.

11. Gordon-Salant S, Fitzgibbons P (2004) Effects of stimulus and noise rate variability on speech perception by younger and older adults. JASA 115: 1808–1817.

12. Cooke M, Lu Y (2010) Spectral and temporal changes to speech produced in the presence of energetic and informational maskers. JASA 128: 2059–2069.

13. Chait M, de Cheveigné A, Poeppel D, Simon J (2010) Neural dynamics of attending and ignoring in human auditory cortex. Neuropsychologia 48: 3262–3271.

14. Luo H, Poeppel D (2007) Phase patterns of neuronal responses reliably discriminate speech in human auditory cortex. Neuron 54: 1001–1010.

15. Ding N, Simon J (2012) Neural coding of continuous speech in auditory cortex during monaural and dichotic listening. J Neurophysiol 107: 78–89.

16. Ding N, Simon J (2013) Adaptive temporal encoding leads to a background-insensitive cortical representation of speech. Journal of Neuroscience 33: 5728–5735.

17. Power A, Foxe J, Forde E, Reilly R, Lalor E (2012) At what time is the cocktail party? A late locus of selective attention to natural speech. European Journal of Neuroscience 35: 1497–1503.

18. Gross J, Hoogenboom N, Thut G, Schyns P, Panzeri S, Belin P, Garrod S (2013) Speech rhythms and multiplexed oscillatory sensory coding in the human brain. PLoS Biology 11: e1001752.

19. Peelle J, Gross J, Davis M (2013) Phase-locked responses to speech in human auditory cortex area enhanced during comprehension. Cerebral Cortex 23: 1378–1387.

20. Millman R, Johnson S, Prendergast G (2015) The role of phase-locking to the temporal envelope of speech in auditory perception and speech intelligibility. Journal of Cognitive Neuroscience 27: 533–545.

21. O’Sullivan J, Power A, Mesgarani N, Rajaram S, Foxe J, Shinn-Cunningham B, Slaney M, Shamma S, Lalor E (2015) Attentional selection in a cocktail party environment can be decoded from single-trial EEG. Cerebral Cortex 25: 1697–1706.

22. Park H, Ince R, Schyns P, Thut G, Gross J (2015) Frontal top-down signals increase coupling of auditory low-frequency oscillations to continuous speech in human listeners. Current Biology 25: 1649–1653.

23. Keitel A, Gross J, Kayser C (2018) Perceptually relevant speech tracking in auditory and motor cortex reflects distinct linguistic features. PLoS Biol 16: e2004473.

24. Zoefel B, Archer-Boyd A, Davis M (2018) Phase entrainment of brain oscillations causally modulates neural responses to intelligible speech. Current Biology 28: 401–408.

25. Schroeder C, Wilson D, Radman T, Scharfman H, Lakatos P (2010) Dynamics of active sensing and perceptual selection. Curr Opin Neurobiol 20: 172–176.

26. Kerlin J, Shahin A, Miller L (2010) Attentional gain control of ongoing cortical speech representations in a “cocktail party”. Journal of Neuroscience 30: 620–628.

27. Horton C, D’Zmura M, Srinivasan R (2013) Suppression of competing speech through entrainment of cortical oscillations. Neurophysiology 109: 3082–3093.

28. Zion Golumbic E, Ding N, Bickel S, Lakatos P, Schevon C, McKhann G, Goodman R, Emerson R, Mehta A, Simon J, Poeppel D, Schroeder C (2013) Mechanisms underlying selective neuronal tracking of attended speech at a “cocktail party”. Neuron 77: 980–991.

29. Ghinst M, Bourguignon M, de Beeck M, Wens V, Marty B, Hassid S, Choufani G, Jousmaki V, Hari R, Bogaert P, Goldman S, De Tiége X (2016) Left superior temporal gyrus is coupled to attended speech in a cocktail-party auditory scene. Journal of Neuroscience 36: 1596–1606.

30. Kong Y, Somarowthu A, Ding N (2015) Effects of spectral degradation on attentional modulation of cortical auditory responses to continuous speech. JARO 16: 783–796.

31. Riecke L, Formisano E, Sorger B, Baskent D, Gaudrain E (2018) Neural entrainment to speech modulates speech intelligibility. Current Biology 28: 161–169.

32. Henry M, Obleser J (2012) Frequency modulation entrains slow neural oscillations and optimizes human listening behavior. PNAS 109: 20095–20100.

33. Henry M, Herrmann B, Obleser J (2014). Entrained neural oscillations in multiple frequency bands comodulate behavior. PNAS 111: 14935–14940.

34. Tal I, Large E, Rabinovitch E, Wei Y, Schroeder C, Poeppel D, Zion Golumbic E (2017) Neural entrainment to the beat: the “missing pulse” phenomenon. Journal of Neuroscience. doi:10.1523/JNEUROSCI.2500-16.2017

35. Tierney A, Kraus N (2014) Neural entrainment to the rhythmic structure of music. Journal of Cognitive Neuroscience 27: 400–408.

36. Doelling K, Poeppel D (2015). Cortical entrainment to music and its modulation by expertise. PNAS 112: E6233–E6242.

37. Meltzer B, Reichenbach C, Braiman C, Schiff N, Hudspeth A, Reichenbach T (2015) The steady-state response of the cerebral cortex to the beat of music reflects both the comprehension of music and attention. Frontiers in Human Neuroscience 9: 436.

38. Harding E, Sammler D, Henry M, Large E, Kotz S (2019) Cortical tracking of rhythm in music and speech. NeuroImage 185: 96–101.

39. Lakatos P, Karmos G, Mehta A, Ulbert I, Schroeder C (2008). Entrainment of neuronal oscillations as a mechanism of attentional selection. Science 320: 110–113.

40. Lakatos P, O’Connell M, Barczak A, Mills A, Javitt D, Schroeder C (2009) The leading sense: supramodal control of neurophysiological context by attention. Neuron 64: 419–430.

41. Lakatos P, Musacchia G, O’Connel M, Falchier A, Javitt D, Schoeder C (2013) The spectrotemporal filter mechanism of auditory selective attention. Neuron 77: 750–761.

42. Lakatos P, Barczak A, Neymotin S, McGinnis T, Ross D, Javitt D, O’Connell M (2016) Global dynamics of selective attention and its lapses in primary auditory cortex. Nature Neuroscience 19: 1707–1719.

43. Besle J, Schevon C, Mehta A, Lakatos P, Goodman R, McKhann G, Emerson R, Schroeder C (2011) Tuning of the human neocortex to the temporal dynamics of attended events. Journal of Neuroscience 31: 3176–3185.

44. Morillon B, Baillet S (2017) Motor origin of temporal predictions in auditory attention. PNAS 114: E8913–E8921.

45. Choi I, Rajaram S, Varghese L, Shinn-Cunningham B (2013) Quantifying attentional modulation of auditory-evoked cortical responses from single-trial electroencephalography. Frontiers in Human Neuroscience 7: 115.

46. Dai L, Best V, Shinn-Cunningham B (2018) Sensorineural hearing loss degrades behavioral and physiological measures of human spatial selective auditory attention. PNAS 115: E3286–E3295.

47. Holt L, Tierney A, Guerra G, Laffere A, Dick F (2018) Dimension-selective attention as a possible driver of dynamic, context-dependent re-weighting in speech processing. Hearing Research 366: 50–64.

48. Rodrigues A, Loureiro M, Caramelli P (2013) Long-term musical training may improve different forms of visual attention ability. Brain and Cognition 82: 229–235.

49. Oxenham A, Fligor B, Mason C, Kidd G (2003) Informational masking and musical training. JASA 114: 1543–1549.

50. Parbery-Clark A, Skoe E, Lam C, Kraus N (2009) Musician enhancement for speech in noise. Ear and Hearing 30: 653–661.

51. Swaminathan J, Mason C, Streeter T, Best V, Kidd G, Patel A (2015) Musical training, individual differences and the cocktail party problem. Scientific Reports 5: 11628.

52. Clayton K, Swaminathan J, Yazdanbakhsh A, Zuk J, Patel A, Kidd G (2016) Executive function, visual attention and the cocktail party problem in musicians and nonmusicians. PLoS ONE 11: e0157638.

53. Slater J, Kraus N (2016) The role of rhythm in perceiving speech in noise: a comparison of percussionists, vocalists and non-musicians. Cognitive Processes 17: 79–87.

54. Deroche M, Limb C, Chatterjee M, Gracco V (2017) Similar abilities of musicians and non-musicians to segregate voices by fundamental frequency. JASA 142: 1739–1755.

55. Meha-Bettison K, Sharma M, Ibrahim R, Vasuki P (2017) Enhanced speech perception in noise and cortical auditory evoked potentials in professional musicians. International Journal of Audiology 57: 40–52.

56. Morse-Fortier C, Parrish M, Baran J, Freyman R (2017) The effects of musical training on speech detection in the presence of informational and energetic masking. Trends in Hearing 21: 1–12.

57. Ruggles D, Freyman R, Oxenham A (2014) Influence of musical training on understanding voiced and whispered speech in noise. PLoS ONE 9: e86980.

58. Madsen S, Whiteford K, Oxenham A (2017) Musicians do not benefit from differences in fundamental frequency when listening to speech in competing speech backgrounds. Scientific Reports 7: 12624.

59. Berens P (2009) CircStat: a Matlab toolbox for circular statistics. J Stat Softw 31: 1–21.

60. Pantev C, Oostenveld R, Engelien A, Ross B, Roberts L, Hoke M (1998) Increased auditory cortical representation in musicians. Nature 392: 811–814.

61. Shahin A, Bosnyak D, Trainor L, Roberts L (2003) Enhancement of neuroplastic P2 and N1c auditory evoked potentials in musicians. Journal of Neuroscience 23: 5545–5552.

62. Schneider P, Sluming V, Roberts N, Bleeck S, Rupp A (2005) Structural, functional, and perceptual differences in Heschl’s gyrus and musical instrument preference. Ann. N.Y. Acad. Sci. 1060: 387–394.

63. Seither-Preisler A, Parncutt R, Schneider P (2014) Size and synchronization of auditory cortex promotes musical, literacy and attentional skills in children. Journal of Neuroscience 34: 10937–10949.

64. Tierney A, Krizman J, Kraus N (2015) Music training alters the course of adolescent auditory development. PNAS 112: 10062–10067.

65. Habibi A, Cahn B, Damasio A, Damasio H (2016) Neural correlates of accelerated auditory processing in children engaged in music training. Developmental Cognitive Neuroscience 21: 1–14.

66. Oostenveld R, Fries P, Maris E, Schoffelen J (2011) FieldTrip: open source software for advanced analysis of MEG, EEG, and invasive electrophysiological data. Computational Intelligence and Neuroscience 2011: 156869.

